# Tellurium Notebooks - An Environment for Dynamical Model Development, Reproducibility, and Reuse

**DOI:** 10.1101/239004

**Authors:** J Kyle Medley, Kiri Choi, Matthias König, Lucian Smith, Stanley Gu, Joseph Hellerstein, Stuart C. Sealfon, Herbert M Sauro

## Abstract

The considerable difficulty encountered in reproducing the results of published dynamical models limits validation, exploration and reuse of this increasingly large biomedical research resource. To address this problem, we have developed Tellurium Notebook, a software system that facilitates building reproducible dynamical models and reusing models by 1) supporting the COMBINE archive format during model development for capturing model information in an exchangeable format and 2) enabling users to easily simulate and edit public COMBINE-compliant models from public repositories to facilitate studying model dynamics, variants and test cases. Tellurium Notebook, a Python–based Jupyter–like environment, is designed to seamlessly inter-operate with these community standards by automating conversion between COMBINE standards formulations and corresponding in–line, human–readable representations. Thus, Tellurium brings to systems biology the strategy used by other literate notebook systems such as Mathematica. These capabilities allow users to edit every aspect of the standards–compliant models and simulations, run the simulations in–line, and re–export to standard formats. We provide several use cases illustrating the advantages of our approach and how it allows development and reuse of models without requiring technical knowledge of standards. Adoption of Tellurium should accelerate model development, reproducibility and reuse.

**Author summary:** There is considerable value to systems and synthetic biology in creating reproducible models. An essential element of reproducibility is the use of community standards, an often challenging undertaking for modelers. This article describes Tellurium Notebook, a tool for developing dynamical models that provides an intuitive approach to building and reusing models built with community standards. Tellurium automates embedding human–readable representations of COMBINE archives in literate coding notebooks, bringing to systems biology this strategy central to other literate notebook systems such as Mathematica. We show that the ability to easily edit this human–readable representation enables users to test models under a variety of conditions, thereby providing a way to create, reuse, and modify standard–encoded models and simulations, regardless of the user’s level of technical knowledge of said standards.

## Introduction

Multiscale dynamical modeling requires the ability to build large, comprehensive, and complex models of biological systems. Examples include the *Mycoplasma genitalium* whole–cell model [1] and the central metabolism of *E. coli* [2]. These models are often composed of many submodels. Typically, submodels are developed and validated by other research teams. Indeed, without the ability to reuse existing models, constructing larger models becomes impractical.

Being able to reuse tools and techniques developed by others is a hallmark of science. Poor reproducibility of biomedical experimental studies has been recognized as a major impediment to scientific progress [3,4]. Much of the focus on poor reproducibility has been on wet lab experiments. However, barriers to reproducibility is also a significant problem in computational studies [5–8]. In recognition of this problem, **reproducibility** has become a central focus of scientific software [9,10]. The general experience of researchers in the field of modeling suggests that a similar problem in poor reproducibility also exists for biomodels. Difficulty in model reproducibility can result from a published model not being deposited in a public repository or from differences in the deposited model and the actual model used for published simulations. In addition, it is difficult for researchers to utilize and modify public models because the standards are not human–readable. This state of affairs imperils continued progress with developing and exploiting biological models.

We propose that reproducible computational studies must satisfy two requirements. First, they must be *transparent*; that is, researchers must be able to inspect and understand the details of the model and the computational experiments. With transparency, researchers can check assumptions and explore variations in computational studies. Second, computational studies must be *exchangeable*; that is, it must be possible for a study done in one computational environment to be done in another computational environment and produce comparable results. For a study to be exchangeable means that other researchers can make use of and build on the published results in their computational environment.

In order to be transparent and exchangeable, a computational model and any simulation experiments must be encoded in a standard format that separates the reusable part of a model and its simulations (i.e. parameters, processes, and kinetics) from the implementation used to simulate it (i.e. the numerical methods and algorithms used to generate results). Models can be described using the Synthetic Biology Markup Language (SBML) [11] or CellML [12] standards. These standards support models based on ordinary differential equations (ODEs), stochastic master equations, and constraint-based modeling [13]. Simulations can be described using the Simulation Experiment Description Markup Language (SED–ML) [14], which encodes the types of simulations, either time-course simulations or steady state computations, that should be run on a model. SED–ML allows specifying the exact numerical algorithms needed to run a simulation using the KiSAO ontology [15], which includes widely used ODE (e.g. LSODA [16,17], CVODE [18]) and stochastic solvers (e.g. Gillespie direct method [19], Gibson algorithm [20]).

In order to facilitate exchanging models and simulations between software tools, SED–ML simulations and SBML / CellML models can be packaged together using COMBINE archives [21]. However, few authoring tools exist for SED–ML and COMBINE archives [22,23]. Furthermore, existing resources require technical knowledge of standards, restricting use of these standards by the modeling community at large. Therefore, an authoring tool is needed that allows a wider range of users to create and edit COMBINE archives containing both models and simulations. We propose that the authoring tool should satisfy five requirements:

1. It should represent the models or simulation specifications in a human–readable form.
2. It should allow the user to easily edit this human–readable representation.
3. It should allow the user to provide narrative, annotations, or comments in order to improve transparency.
4. It should translate the specifications into an implementation that can be used to run simulations.
5. It must be capable of repackaging the model and/or simulation in a standard form that is usable by other tools.

To address these requirements, we have developed the Tellurium Notebook environment, which extends the literate notebook concept used by tools like Jupyter [24] and Mathematica [25] to support community standards in systems biology. Whereas Jupyter notebooks contain code and narrative cells, Tellurium adds a third cell type for representing models and simulations encoded as standards. Our tool allows modeling studies to be constructed in a notebook environment and exported using community standards. This workflow provides both transparency, through a literate notebook, and exchangeability, through seamless, fluid support for standards.

Tellurium supports embedding human–readable representations of SBML [26] and SED–ML [27] directly in cells. These cells can be exported as COMBINE archives which are readable by other tools. We refer to this human–readable representation as *inline OMEX* (after Open Modeling and EXchange, the encoding standard used by COMBINE archives). Inline OMEX cells operate in much the same way as code cells, i.e. they have syntax highlighting and are executable. Executing an inline OMEX cell runs all SED–ML simulations in the cell, producing any plots or reports declared in the SED–ML. A major advantage of this approach is that it offers a means of authoring transparent, exchangeable modeling studies without requiring technical knowledge of standards.

## Results

We demonstrate the benefits for reproducibility provided by Tellurium Notebook with case studies. In the first case study, we explore the impact of variations in the value of a parameter in a model of yeast. Such explorations are frequently done to determine if a model is applicable to conditions beyond those in the original model, an important consideration for testing model validity. The second case study evaluates if a model implementation produces results that are comparable to those in the original study via a series of tests which cover important dynamical properties of the model.

## Case Study 1: Assessing Previously Unexplored Model Parameters

In order to meet our requirements for reproducibility, it is not sufficient to simply recreate a simulation. Rigorous reproducibility requires the ability to reuse, expand, and test existing models under a variety of circumstances. This first case study shows how Tellurium facilitates conducting new experiments with an existing model, including the packaging of the model and the experiments as a COMBINE archive. The study is based on a model of autonomous metabolic oscillations in yeast and associated numerical studies [28]. The model has a cooperativity parameter *m* for which no specific justification is provided. We explore a range of values of *m* to understand how this parameter influences the dynamics in the model.

When grown in continuous culture, yeast exhibit oscillations that can be maintained for months and can cause the temporal separation of many cellular functions in a synchronized way [29]. This type of dynamics belongs to a broader class of coupled oscillators that are thought to be highly important in the organization of circadian rhythms and play a role in the regulation of diurnal physiological activity in various species [30,31]. Wolf, et al. [28] developed a model that uses metabolic coupling mediated in part by the *H*_2_*S* pathway, one of the known contributors to respiratory oscillations in yeast [29], to explain experimentally observed synchronized oscillations in yeast cultures.

The model contains 21 reactions and is available via the BioModels repository as `BI0MD0000000090` [32]. The reaction `v11a` is a key part of the *H*_2_*S* pathway. As shown in Fig 1, standard–encoded SBML for this reaction is difficult to decipher, much less comprehend. As with other authoring tools, Tellurium provides a concise, human–readable abstraction of this encoding. However, unlike other tools, this abstraction covers the entire SBML model and even covers entire COMBINE archives, as we show shortly. We focus on the human readable representation of the reaction `v11a`, including the reaction kinetics using the Hill coefficient *m*.

**Fig 1.**
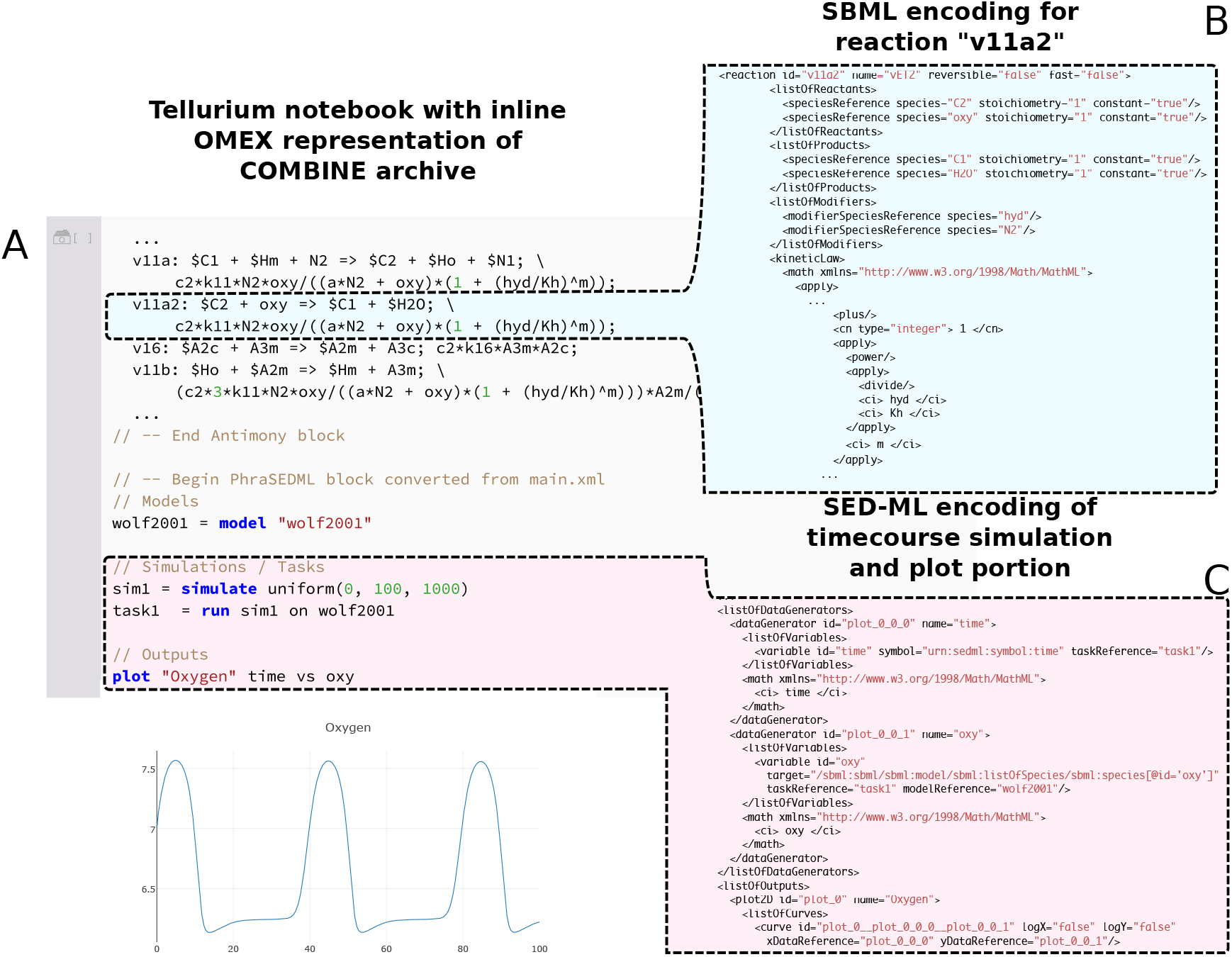
A comparison of Tellurium’s human–readable representation of a COMBINE archive shown in a Tellurium notebook (A) and excerpts from the equivalent SBML (B) and SED–ML (C) encodings. Tellurium’s in-line OMEX format contains human-readable representations of both SBML and SED–ML (A). (B) contains the SBML encoding for a single reaction. The single–line human–readable form of this reaction is highlighted in part (A) for comparison. Using the SBML encoding, it is difficult to modify the reaction stoichiometry or kinetic law, whereas this task is easy in Tellurium. This example makes use of a model of respiratory oscillations in yeast [28] and is available as a COMBINE archive [33]. Note, when using Tellurium, only the human–readable representation is displayed and can be modified by the user to automatically generate modifications in the hidden underlying standard representations.

Clearly, readability is essential for model transparency. However, readability is essential for model reuse as well. To demonstrate this, we convert this SBML–only model into a COMBINE archive containing both SBML portions describing the model and SED–ML portions describing the simulation. We then show how Tellurium’s human-readable format permits easy modification of the published model and simulations contained in the COMBINE archive.

In order to create a SED–ML specification for this model, we need to define four steps in the workflow, which correspond to distinct elements in SED–ML: (1) model definition, (2) simulation, (3) task specification, and (4) output generation. For (1), models can be defined in Tellurium’s human–readable format by referencing SBML or CellML files in the same COMBINE archive, with the option of including parameter replacements. For (2), SED–ML simulations can be either timecourse simulations or steady state computation, and can reference a specific algorithm (e.g. LSODA), or a generic implementation using the KiSAO ontology [15]. Tellurium uses predefined keywords such as lsoda (an ODE solver implementation [34]) to refer to popular implementations. In SED–ML, simulations are specified independently from models. This allows model and simulation elements to be reused in different combinations. For (3), SED–ML uses task elements to describe these combinations. Finally, the output elements of (4) can be plots or reports and allow users to access the output of tasks. Tellurium’s human–readable format allows defining a SED–ML model by instantiating the same SBML model with different parameter values (m in this example) using the syntax:

~~~
mymodel = model “wolf2001” with paraml=value2, param2=value2 …
~~~

with the param/value pairs being replaced by the corresponding parameter ids and values respectively. We use this syntax to instantiate five copies of the model and explore the values *m* =1, 2, 4, 8, 16. Since we do not know *a priori* if the value of *m* affects the timescale of the dynamics, we also create two simulations using different durations. Finally, we create a task for each model/simulation combination and plot the results on their respective timescales. Fig 2 shows that the value of *m* drastically affects the dynamical behavior of the system, abolishing the periodicity of the oscillations at *m* = 8 and ceasing them entirely at *m* = 16. Smaller values of *m* also affect the phase and amplitude of the oscillations.

**Fig 2.**
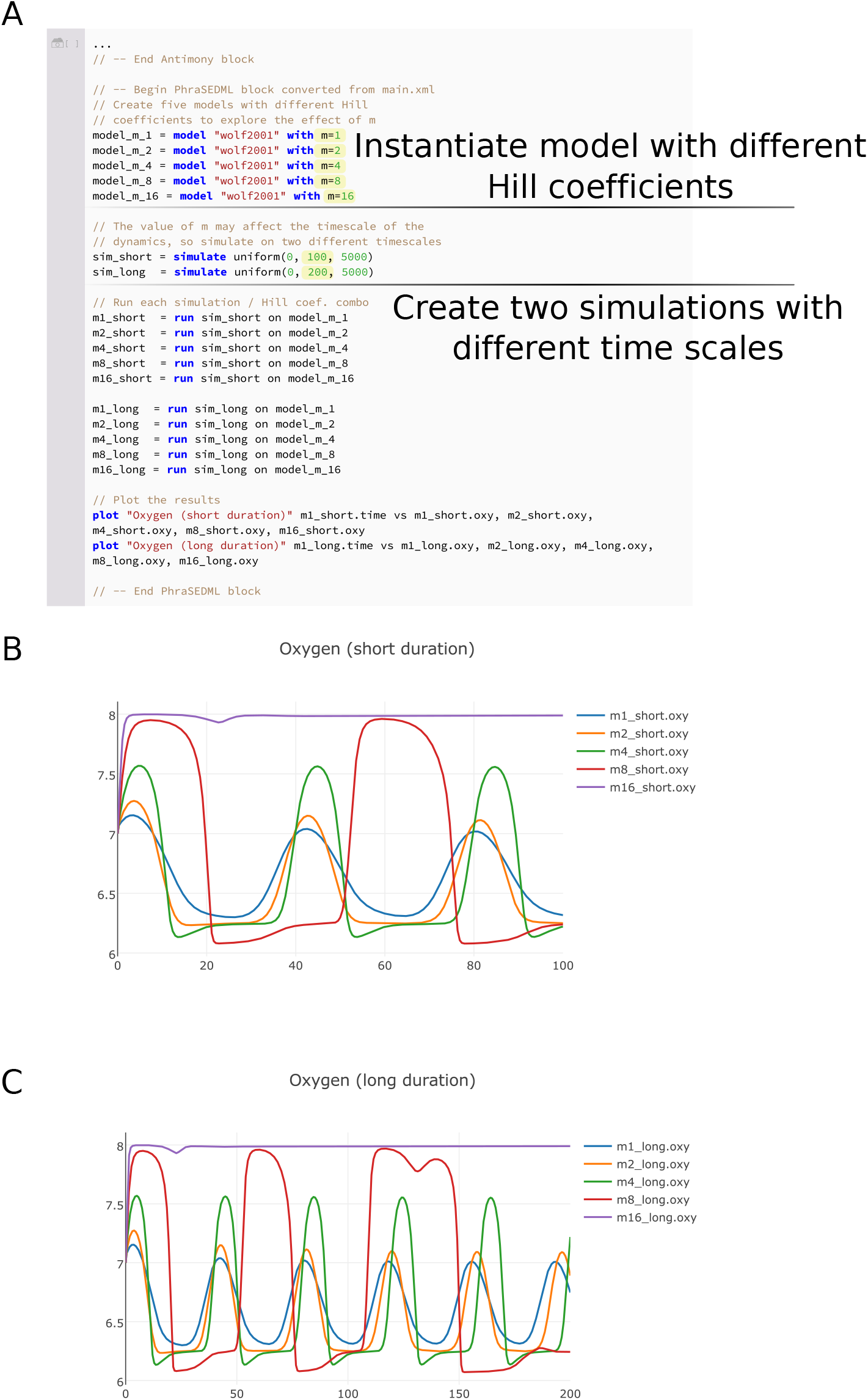
An example of using Tellurium to edit the respiratory oscillations model introduced in Fig 1. To investigate the effect of the Hill coefficient *m*, we used Tellurium’s human-readable representation of SED–ML to create five instantiations of the model using values of 1, 2, 4, 8, and 16 for *m* (A). We then simulated each of these instantiations on two different timescales and plotted the respective results for short (B) and long (C) simulations. Tellurium’s COMBINE archive support allows this model and simulation to be exported to other tools, as shown in Fig S1

This case study shows that Tellurium provides an efficient means of converting SBML models into exchangeable COMBINE archives containing simulation components. Furthermore, COMBINE archives can contain important dynamical information about the model, such as the influence of the parameter *m* that we explored in this study. In order to demonstrate exchangeability of this study, we have exported it to the SED-ML Web Tools [22] (Fig S1) and iBioSim [35] (Fig S2).

## Case Study 2: Reproducibility Through In–depth Variational Studies

Reproducibility requires that a model implementation produces results consistent with the original study, especially if a different authoring tool is used. In order to provide criteria for judging whether a model reproduction is consistent with the original, a set of testing criteria are required, similar to the concept of unit testing in software. However, researchers seldom perform extensive checks on the dynamics of models before using them. This is due in part to the lack of tool support for easily modifying and producing variants of models and simulations encoded in exchangeable formats. Tellurium’s authoring features enable modelers to encode dynamical unit tests in COMBINE archives, thereby providing a way to verify that a model has been correctly reproduced.

For this case study, we reproduce a highly–detailed model of syncytial nuclear divisions in the *Drosophila* embryo [36] through testing the model’s dynamics under different conditions. In many insect species, the embryo enters a period of rapid mitotic division without cytokinesis [37] immediately following fertilization. In *Drosophila,* 13 of these divisions occur within 3 hours of fertilization [36]. These divisions are regulated by metaphase promoting factor (MPF), a complex between cyclin (specifically the cyclin CycB in this model) and cyclin–dependent kinases (Cdk). CycB subunits tend to be the limiting factor in complex formation, and are thought to regulate mitotic division. CycB availability is controlled by the anaphase promoting complex (APC), which targets CycB for degradation. However, in *Drosophila,* the levels of CycB appear to remain high during the first 8 mitotic divisions [38]. This observation can be reconciled with known mechanisms by assuming that CycB degradation only occurs in the vicinity of the mitotic spindle [36,39,40], despite the absence of a nuclear envelope during the mitotic divisions. To account for this hypothetical local degradation of CycB, the model artificially separates the cytoplasm into two “compartments,” with a cytoplasmic compartment representing the cell and a nuclear compartment representing the volume in the vicinity of the mitotic spindle.

As a starting point, we use the COMBINE archive encoding of this model by Scharm and Tourè [41]. This archive contains SBML derived from biomodel `BI0MD0000000144`^1^, which is intended to reproduce Fig 1 of [36]. However, the archive does not contain more extensive tests of the model’s dynamics, such as whether the model can be used to reproduce several other simulations described in the paper. The initial variant encoded by the COMBINE archive and shown in Fig 3B and C is based on a model with a constant level of the phosphatase String, whereas in reality String levels change over the course of the mitotic cycles. String regulates MPF via a positive feedback loop, and has been shown to peak at the seventh or eighth cycle of the mitotic divisions [36]. To account for this, Calzone et al. [36] posited that String mRNA is degraded by a hypothetical factor “X,” causing the synthesis rate of String to drop over time. Therefore, we have modified the SED–ML of the original COMBINE archive [41] as follows to include the synthesis and degradation of String. We are able to reproduce Fig 3 of [36] by making these modifications to the original COMBINE archive:

**Fig 3.**
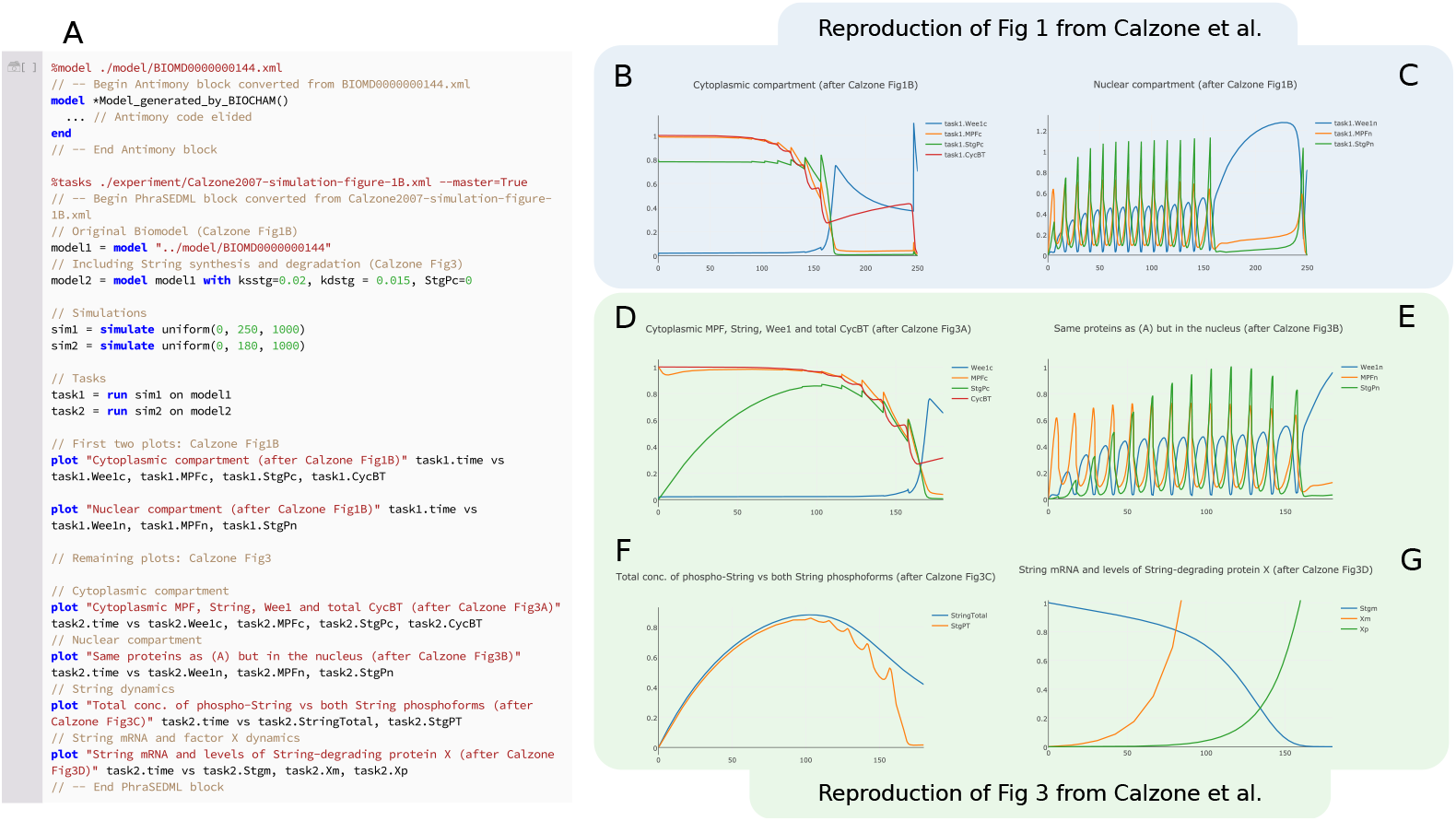
Using Tellurium to reproduce model variants in [36] and repackage as a COMBINE archive. To demonstrate the use of COMBINE archives for encoding model variants, we began with a COMBINE archive describing a single variant of this model without String synthesis or degradation [41], which reproduces Fig 1B of [36] (plots B and C here). We then used Tellurium to add a variant describing String degradation, which reproduces Fig 3 of [36] (plots D through G here). Plots B and D show the cytoplasmic compartment of the model. Plots C and E show the nuclear compartment (defined as the spatial region around the mitotic spindle). Plot F shows the levels of total String and its phosphorylated state. Plot G shows the level of String mRNA and protein factor X, which degrades String mRNA. Note the y–axis scale on plot G was manually adjusted to show the mRNA dynamics. The subplots in this figure intentionally have different durations, after Calzone et al [36]. The model in [36] was authored using BIOCHAM [42]. Our model reproductions that reproduce these plots are available as a COMBINE archive [43].

- Enable synthesis and degradation of String by setting the parameters `ksstg=0.02` and `kdstg=0.015` respectively.
- Set the initial concentration of total String to zero by setting `StgPc=0`.
- Compute the total amount of unphosphorylated String by adding the rule `StgT:= (1 – N*E_1)*Stgc + N*E_1*Stgn`.
- Compute the total amount of String in the cell by adding the rule `StringTotal:= StgPT + StgT`.

Tellurium makes it easy to encode both the original variant, without String synthesis and degradation, and the variant including these terms in a COMBINE archive [43]. Fig 3 shows the results of executing this COMBINE archive in Tellurium. We have thus expanded the dynamical test cases for this model, as it now reproduces two simulations from two different variants described by the original authors (Fig 1 and 3 of [36]), enabling better coverage of the model’s dynamics.

In order to gain insight into the regulatory mechanism controlling the mitotic divisions, and understand the transitions that control the exact number of these divisions, Calzone et al. performed a one–parameter bifurcation analysis [36]. Bifurcation analyses probes the number and position of steady states and other types of attractors as a function of a parameter. The oscillations shown in Fig 3 are the result of discrete division events, and the behavior shown does not represent a limit cycle. However, the model can be shown to exhibit limit cycle behavior by 1) removing all discrete events and 2) fixing the number of divisions by introducing the variable *C* as a cycle counter. The number of nuclear compartments is then given by *N* = 1.95^*C*^ (1.95 is a scaling factor described in [36]). For a given cycle number *C*, MPF exhibits limit cycle oscillations, although the amplitude and period of these oscillations changes with the cycle number. At low cycle number, Calzone et al. observed that these oscillations were dominated by the negative feedback effect of cyclin degradation, whereas for large cycle number (*C* ≥ 12), positive feedback via control of phosphorylated MPF by the kinase Wee and phosphatase String contributes to the oscillations.

SED–ML does not support bifurcation analysis, precluding us from reproducing that part of the study in an exchangeable format. However, it is still possible to test the change in regulatory shift from negative to positive feedback. Instead of a bifurcation diagram, we compare the limit cycle behavior of the original model to a model variant with reduced Wee and String activation and deactivation rates. This slows the timescale of the positive feedback component of the model. Fig 4 compares the behavior of the original model at early and late cycle numbers with the variant containing attenuated positive feedback. Whereas the normal model exhibits stable limit cycle oscillations at both *C* = 1 and *C* = 12, the oscillations in the attenuated model are transient at late cycles (*C* = 12) but not at early cycles (*C* = 1). This observation suggests that String and Wee dynamics are indeed crucially important for late cycle oscillations, but not for early cycle oscillations, confirming the shift in regulatory mechanism. These simulation thus form a third set of unit tests for the model, encoded as a COMBINE archive [44].

**Fig 4.**
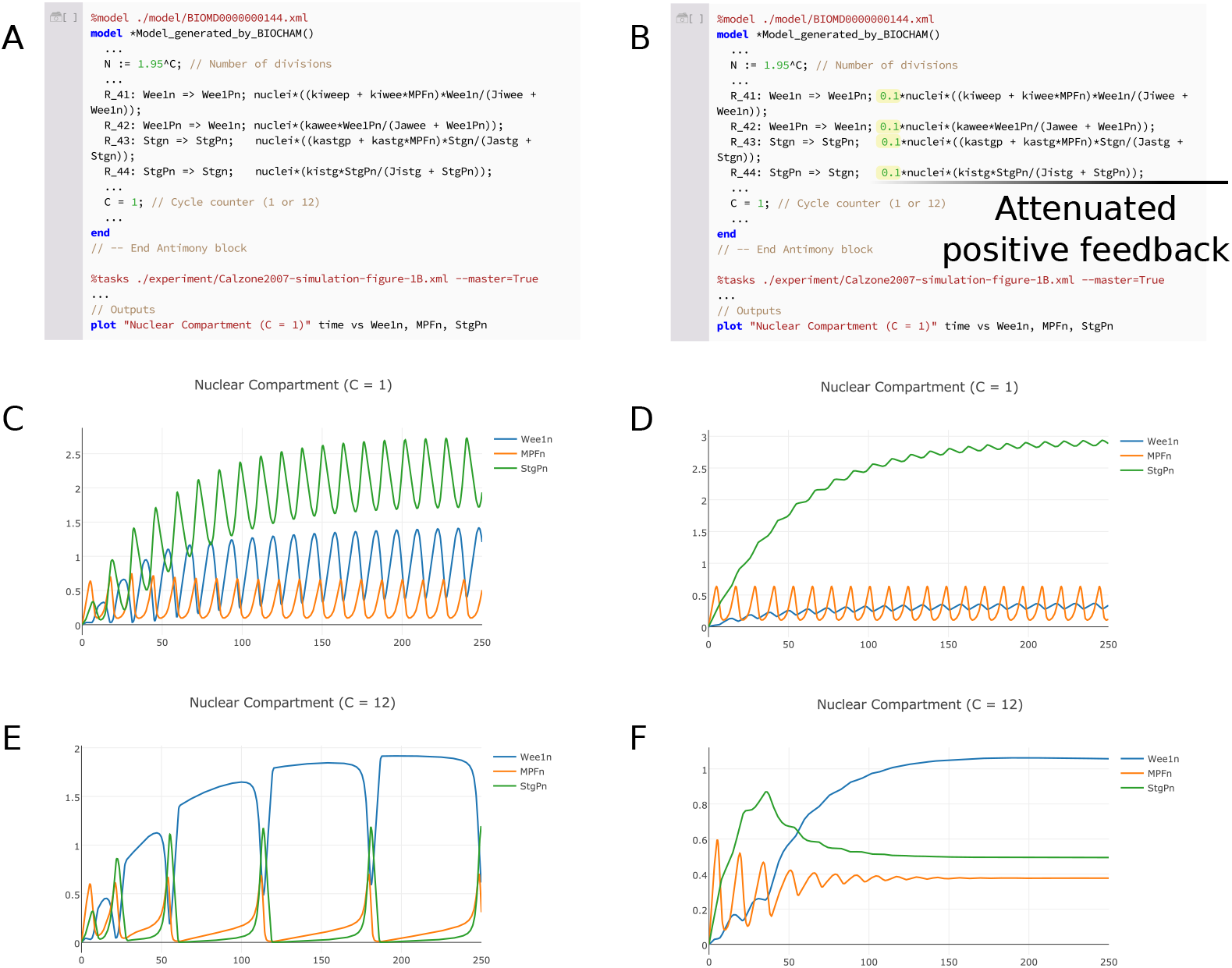
Testing the shift in regulatory mechanism of mitotic oscillations. To verify the observation [36] that the number of mitotic divisions in the *Drosophila* embryo is governed by a shift from negative to positive feedback, we first removed all discrete events and introduced the variable *C* such that *N* = 1.95^*C*^. We then compared the limit cycles produced by this eventless model (left) with those produced by a variant with attenuated positive feedback from the regulators Wee and String (right). Attenuation was achieved by decreasing the rates of the phosphorylation and dephosphorylation of Wee and String. The original model exhibits stable limit cycle oscillations for both early cycles (C), which are putatively dominated by negative feedback, and late cycles (E), which are putatively dominated by positive feedback. The attenuated model only exhibits stable oscillations at early cycles (D), suggesting that positive feedback does indeed play a role in late cycle oscillations (F). Our model reuse and modification study is available as a COMBINE archive that reproduces the figure shown and facilitates further modification and reuse [44].

In summary, using Tellurium’s editing capabilities, we have created an extensive set of unit tests for dynamical behavior this model, which we exported as a COMBINE archive and imported into another tool as shown in Fig S3. Creating these tests required a means of quickly editing and expanding upon both the SBML and SED–ML embedded in the COMBINE archive. Tellurium’s notebook approach allows us to satisfy these requirements, and provides an integrated workflow for testing the dynamical behavior of the model.

### Interoperability Concerns & Test Cases

In order to achieve the exchangeability requirement of reproducibility, broad standards compliance is necessary. A small number of test cases, such as the first two case studies, is not sufficient to ensure interoperability with other software. During Tellurium’s development, we gathered a number of COMBINE archive exemplars from the literature, other software tools, and our own archives. We have provided these archives as a resource to other developers by making them publicly available online. The test archives are structured to separate examples with advanced SED–ML features from those with basic SED–ML usage, enabling tool developers to implement incremental support for the standard. Table S1 lists all test archives and how to obtain them.

### Advanced SED—ML Support

In order to address the requirement of broad standards compliance, we tested Tellurium against a set of tests provided by the SED–ML Web Tools [22]. These tests utilize advanced features of the SED–ML standard, and are designed to demonstrate the standard’s coverage of different types of analysis. Table S2 lists all files used in this test set, and Fig S6 shows the results of exporting these files to Tellurium and back again.

## SBML Test Suite COMBINE Archives

The SBML Test Suite [45] is a collection of dynamical models along with expected trajectories designed to test software tools for compliance with the SBML standard. Each test case contains a SBML model, simulation parameters encoded in SED–ML, expected trajectories encoded as a common–separated values (CSV) file, and graphical plots for reference. We converted each of these 1196 test cases into COMBINE archives containing the SBML models, SED–ML simulations, and CSV expected results and used these COMBINE archives as a benchmark for Tellurium’s support for standards. The results of this benchmark are shown in Table S3.

## Discussion

In order for the conclusions of a research study to be valid, the models used in the study must be reliable. Using SED–ML to reproduce the dynamics of a model and compare these dynamics with expected values adds crucial value to the integrity and validity of studies that reuse or expand on the model. As an exchangeable format, SED–ML is confined to the intersection of the most common features available in dynamical modeling tools, which leaves out certain useful types of analysis (e.g. bifurcation analysis). However, we argue that the use case of SED–ML is not to serve as a replacement for current analysis methods. Instead, SED–ML is a tool to test the dynamical behavior of models before using them. For example, while we were not able to reproduce the bifurcation analysis of the mitotic division study [36] in an exchangeable format, we were able to verify the observations regarding the shift in regulatory mechanism, and in doing so gained new insight from this alternative approach. A researcher may also wish to verify that the model reproduces certain expected behaviors. For example, if the model is expected to exhibit switch–like behavior, does this behavior occur at the correct input threshold? For models with feedback, such as integral feedback control [46], does the output exhibit robustness in the presence of perturbations? These types of validation require expert knowledge of the system. While there are tools and resources to help with this, the most important point for conveying this information to other researchers is to encode it as transparently and lucidly as possible, which is achieved using the literate notebook approach described here.

Tellurium’s approach of blending standards with literate coding enables researchers to create rich, detailed workflows incorporating community standards. Tellurium allows the models and simulations from these notebooks to be shared with other tools via COMBINE archives. This allows other users to import these models and simulations and reproduce them using independently developed software tools. This is consistent with our original definition of reproducibility, as it enables robust cross–validation of results between tools, as opposed to simply repeating a previous simulation. It also helps ensure that the tools themselves are accurate and free of idiosyncrasies that could affect the analysis results. Model repositories such as BioModels [47,48], JWS Online [49], and the CellML model repository [50] have enabled widespread support for the SBML and CellML standards. We believe that better tool support for SED–ML and COMBINE archives will help create a trend toward better adoption of these formats by repositories.

### Comparison with Existing Software

Many dynamical modeling tools support exchanging models via the SBML format, including COPASI [51,52], SBW [53], iBioSim [35], PathwayDesigner [54], CellDesigner [55,56], VCell [57-59], CompuCell3D [60], PySCeS [61], BioNetGen [62], and PySB [63]. These tools have diverse feature sets and intended use cases, such as tissue modeling (CompuCell3D), rule–based modeling of molecular complexes (BioNetGen, PySB, VCell), and general modeling and simulation (all others). The tools also have different forms of user interaction, such as graphical user interfaces (COPASI, iBioSim, VCell) and graph–based network editors (CellDesigner, PathwayDesigner). Python–based tools such as PySCeS [61] and PySB [63] can be used with a Jupyter notebook, but do not feature integration of standards with the notebook itself. In general, Tellurium is useful when the user wishes to interactively edit and test standard–encoded models and simulations or produce presentations and PDFs of modeling studies.

Tellurium’s Python foundation makes it easy to combine with other Python–based software such as PySCeS, COBRApy [64], and PySB. There are also many specialized Python packages for specific tasks such as moment closure approximation for stochastic models [65], parameter estimation [66], Bayesian inference [67], and estimating rate laws and their parameter values [68].

In biomedical research, certain tools have been created specifically to facilitate reproducible research. One such tool is Galaxy [69]. Galaxy is a web–based tool which allows users to create workflows describing experiments, e.g. metagenomic studies [70]. A similar tool with a focus on web services and which supports SBML–based workflows is Taverna [71]. Galaxy and Taverna allow users to annotate each step of the workflow, which provides a way for others to follow and understand the chain of reasoning used in the workflow’s construction. This satisfies the requirement of transparency, as it allows users to view the sequence of steps used to produce a result. Although this approach is very different from a literate notebook in terms of the way the user interacts with the system, it shares the goal of allowing the user to see the sequence of steps used to produce a result and interrogate the specific procedure used in each of the steps. Galaxy and Taverna also allow users to share workflows via the web. However, neither tool attempts to directly address the problem of exchangeability with other software tools.

VisTrails [72] is another workflow system based on visual design. VisTrails focuses primarily on generating rich, three–dimensional diagrams and visualizations based on input data and a specific sequence of steps. VisTrails also saves all changes made to a workflow and allows users to view previous versions, a concept termed “retrospective provenance” [73]. However, this approach also lacks exchangeability. Furthermore, while graphical tools may be more accessible because they abstract away the underlying algorithms, it can be difficult to isolate and correct software errors when a step fails due to bad input or an internal error.

Many other research software systems make use of notebooks, and some incorporate special extensions. StochSS [74], the GenePattern Notebook [75], the SAGE math system [76], and the commercial Mathematica software [25] all utilize notebooks which are specially tailored or feature special extensions for each respective application. However, none of these approaches attempt to solve the problem we address: workflow integration with exchangeable standards. Our usage of the literate notebook approach is intended to satisfy two specific requirements, which are distinct from other use cases: 1) to make these standards easy for humans to read, understand, and modify, without requiring expert knowledge of the technical specifications of the standards, and 2) provide an integrated workflow which facilitates exchangeability with other software.

The notebook approach used by Tellurium also has disadvantages. For example, it is difficult to use notebooks with a version control system in a meaningful way. Furthermore, large or complex analyses can be difficult to orchestrate using notebooks, as interacting with a large notebook with many cells can be cumbersome. Nevertheless, we believe that Tellurium’s approach is highly useful in many crucial use cases, including testing models, experimenting with model variants, and as a final step in producing an analysis for other researchers in a transparent, visual presentation.

## Conclusion

In order to build larger, more complete, and more accurate dynamical models of cells and tissues, it will be necessary to reuse models of subsystems. This is currently very difficult due to the time–consuming and laborious process of manually reconstructing models from the literature, or manually verifying third–party SBML models. Tellurium provides support for encapsulating both a model and its dynamics in a community–developed standard format, the COMBINE archive. This archive can contain the model as well as a number of simulations which test various dynamical properties of the model. Tellurium allows users to create COMBINE archives easily from SBML models, or import and modify preexisting COMBINE archives.

Tellurium integrates SBML, SED–ML, and COMBINE archives within a notebook environment, making it exceptionally easy for users to work with these standards, and obviating the need for users to understand the technical specifications of the standards. The availability of authoring tools such as Tellurium will make it possible for model repositories to begin implementing support for SED–ML and COMBINE archives. Indeed, the JWS Online repository [49] already has support for exporting COMBINE archives of models and simulations, which can be read by Tellurium. We hope that other databases will follow suit so that it will be possible to automatically extract dynamical information from these repositories.

Tellurium’s human–readable representation of COMBINE archives is highly important for facilitating model modification as we describe here. This feature enables researchers to experiment with models using alternate parameterizations in order to test the dynamical behavior of the models under varying conditions. We hope that this will lead to more robust models which lead to biological insight by providing predictions under a wide range of circumstances, as with the case studies presented here.

## Future Work

There is a clear need to support exchangeability of simulation experiments in order to allow researchers to build larger, better tested, and more comprehensive models. Tellurium’s built-in support for exchangeability comes from the SBML and SED–ML standards. This allows Tellurium to support the widest possible range of software tools, but also prevents exchanging studies not covered by SED–ML’s vocabulary of predefined simulation types. Due to delays associated with standardizing and implementing features, SED–ML tends to lag several years behind other systems which do not rely on standardization. Thus, SED–ML has the advantage of stable support from a wide range of tools, but has the disadvantage of lacking the flexibility to encode custom studies based on recent advancements in model simulation.

In order to provide a more flexible platform for encoding simulation studies, new solutions are need. One such solution would be to extend SED–ML with generic scripting capabilities. Another solution would be to build an alternative platform for exchanging simulation experiments. For example, the SESSL [77] software tool also provides a means for encoding and exchanging simulations. Whereas SED–ML uses a standardized XML schema to describe simulations, SESSL uses a domain-specific-language implemented using the Scala programming language. This allows users to mix in Scala code to access features not yet available via SESSL’s public interface. However, this approach is not language–agnostic and is tied to Scala and its low–level execution engine. The SED–ML standard, in contrast, does not constrain the low–level operation of its implementations.

In this paper, we have argued for modelers to construct “unit tests” for dynamical models by including model variants as in the study by Calzone et al. [36]. We have shown that these variants are easy to construct and encode in COMBINE archives using Tellurium, but we have not addressed how to validate these tests in an automated way. Due to simulation algorithm differences between tools and the presence of multiple steady states in some models, performing a direct numeric comparison between steady state values or timecourse traces may be too fragile to be useful.

The BIOCHAM software tool [42] employs an interesting solution by using temporal logic constructs to make assertions about properties of model timecourse dynamics. Using this approach, it would be possible, for instance, to make semi-quantitative assertions such as “species X exhibits oscillations with a period of 100 ± 50*mHz*”. These logical constructs could be used in lieu of a direct numerical comparison to validate the dynamics of a model. A practical solution to the problem of validating model timecourse dynamics would likely make use of semi-quantitative assertions such as “Is the number of oscillations of X at least 10,” “Does Y exhibit a peak value of at least 100 nM,” or “Does the response time of the system fall within a certain range?”

However, we believe that several important questions remain before such a validation method will be useful in practical contexts, such as what is the minimal set of formal logic expressions sufficient to capture any useful assertions, and what are the best practices for encoding assertions? For example, should the assertions strive to use relative relationships between model quantities, such that reparameterizing the model does not affect the assertions, or should they be valid only for a single given parameterization? In the former case, how should model variations be generated to test assertions? We believe that implementing automated testing of dynamical models requires addressing these questions in a well thought–out way. Until then, we believe that manually comparing results encoded as COMBINE archives as in the studies presented here will provide immediate benefits to reproducibility. For moderate–size models such as the Calzone study, we have shown that this approach is a practical solution.

## Availability

Tellurium Notebook is available as a standalone app (`tellurium.analogmachine.org`) or as a collection of Python packages hosted on the Python Package Index (`pypi.python.org`) for 64-bit versions of Mac OS X, Windows, and Linux. The Tellurium Python packages support Python 2.7, 3.4, 3.5, and 3.6. The notebook app comes bundled with Python 3.6 and all requisite packages. The source code of Tellurium (github.com/sys-bio/tellurium) is licensed under the Apache license, version 2.0. Tellurium incorporates or makes use of other software, such as *nteract, Plotly* (http://plot.ly), *Python, libSBML, libSEDML,* and others, which are licensed under their respective terms. See tellurium.analogmachine.org for links to installation instructions, documentation (`tellurium.readthedocs.io`), and the source code (github.com/sys-bio/tellurium).

## Acknowledgments

JKM, KC, LS, SG, and HMS were supported by NIH grants GM081070-01, GM123032-01A1, NHLBI U01HL122199-02. JH is supported by the Moore/Sloan Data Science Environments Project at the University of Washington supported by grants from the Gordon and Betty Moore Foundation (Award #3835) and the Alfred P. Sloan Foundation (Award #2013-10-29). MK is supported by the Federal Ministry of Education and Research (BMBF, Germany) within the research network Systems Medicine of the Liver (LiSyM, grant number 031L0054). SG was supported by NIAID Modeling Immunity for Biodefense HHSN266200500021C. SCS was supported by NIH grant U19 AI117873. The content is solely the responsibility of the authors and does not necessarily represent the views of the National Institutes of Health, the Betty Moore Foundation, or the Alfred P. Sloan Foundation. We wish to thank Frank Bergmann and Chris Myers for their help and guidance in diagnosing and fixing interoperability problems.

## Supporting information

All COMBINE archives used in this paper can be obtained at https://github.com/0u812/tellurium-combine-archive-test-cases.

---

**Fig S1.**
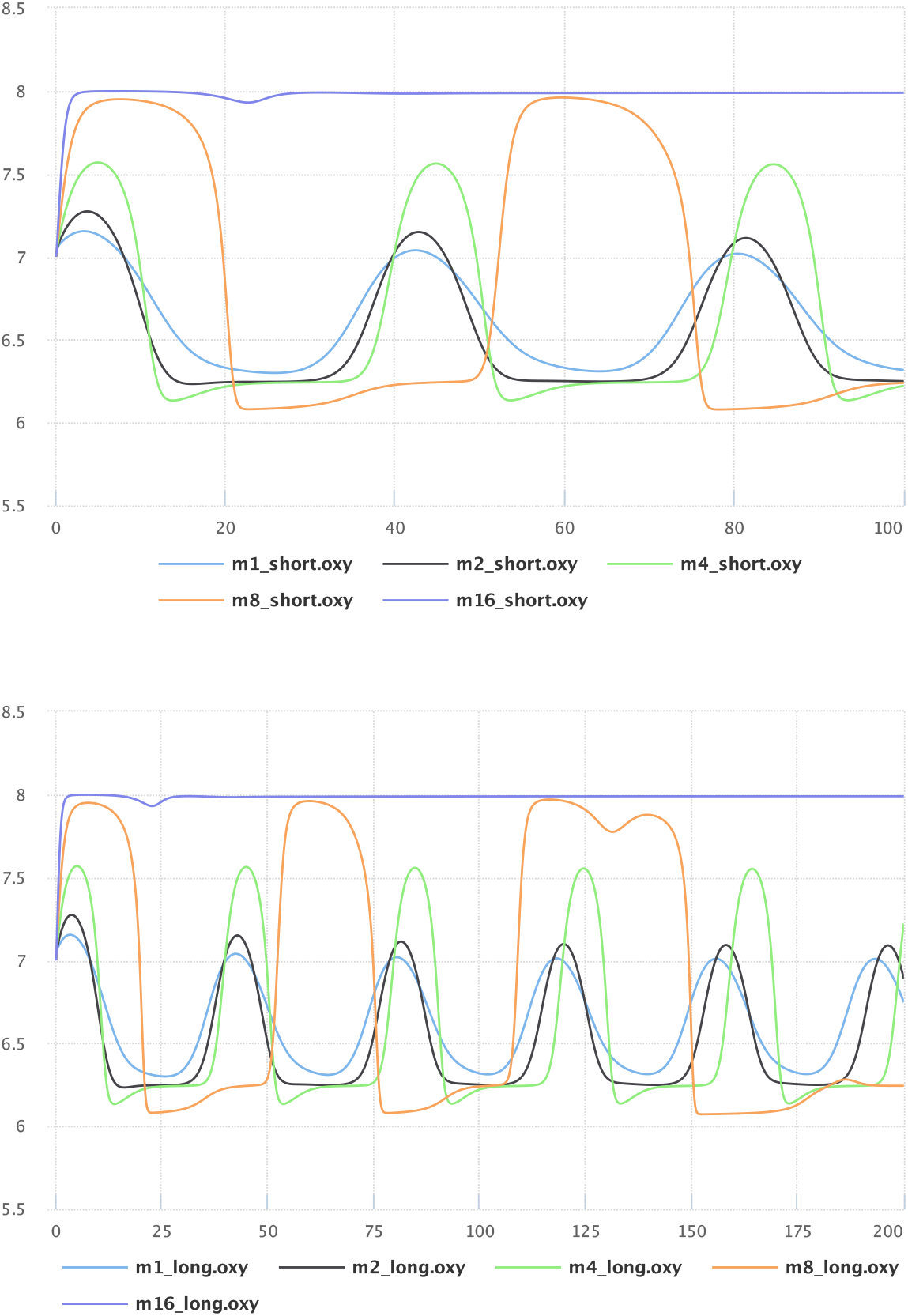
Demonstrating exchangeability of COMBINE archives containing SBML and SED–ML. The respiratory oscillation case study in Fig 2 was exported to a COMBINE archive from Tellurium, imported into the SED–ML Web Tools [22], and used to generate plots to verify that the simulation results were identical to Fig 2. The COMBINE archive used to create this example can be found online [78].

**Fig S2.**
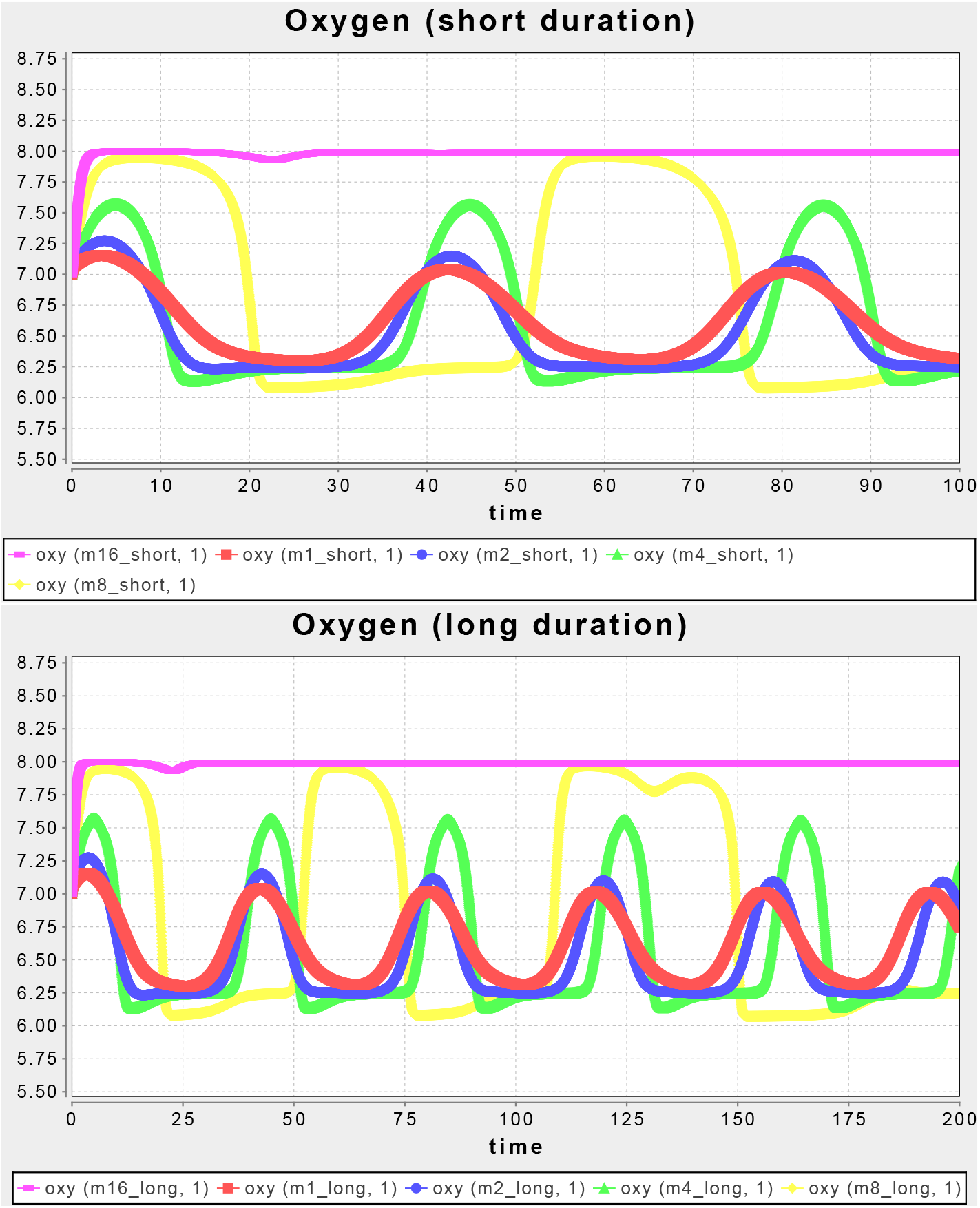
A second test of exchangeability. To demonstrate exchangeability between multiple tools, the same Hill coefficient case study as shown in Fig S1 was exported to iBioSim [35] and used to produce identical plots. This shows that COMBINE archives are sufficiently flexible to be exchanged between different tools, despite the limited number of tools which currently support the format.

**Fig S3.**
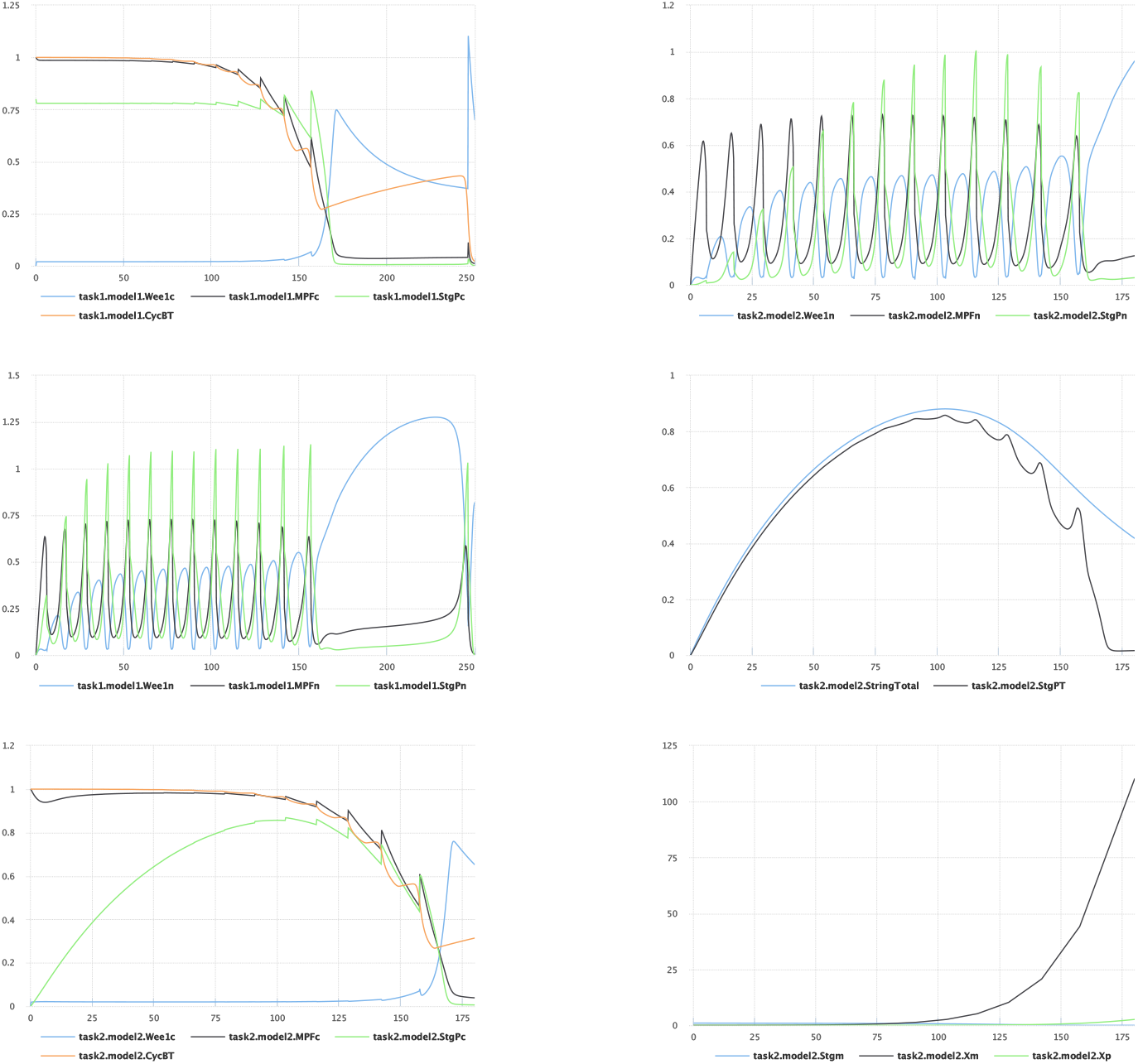
Demonstrating exchangeability of the second case study. To show that the extended set of simulations from Fig 3 can be exchanged with other tools via a COMBINE archive, we exported the study in Fig 3 to the SED–ML Web Tools and verified that the plots were identical to Fig 3.

**Fig S4.**
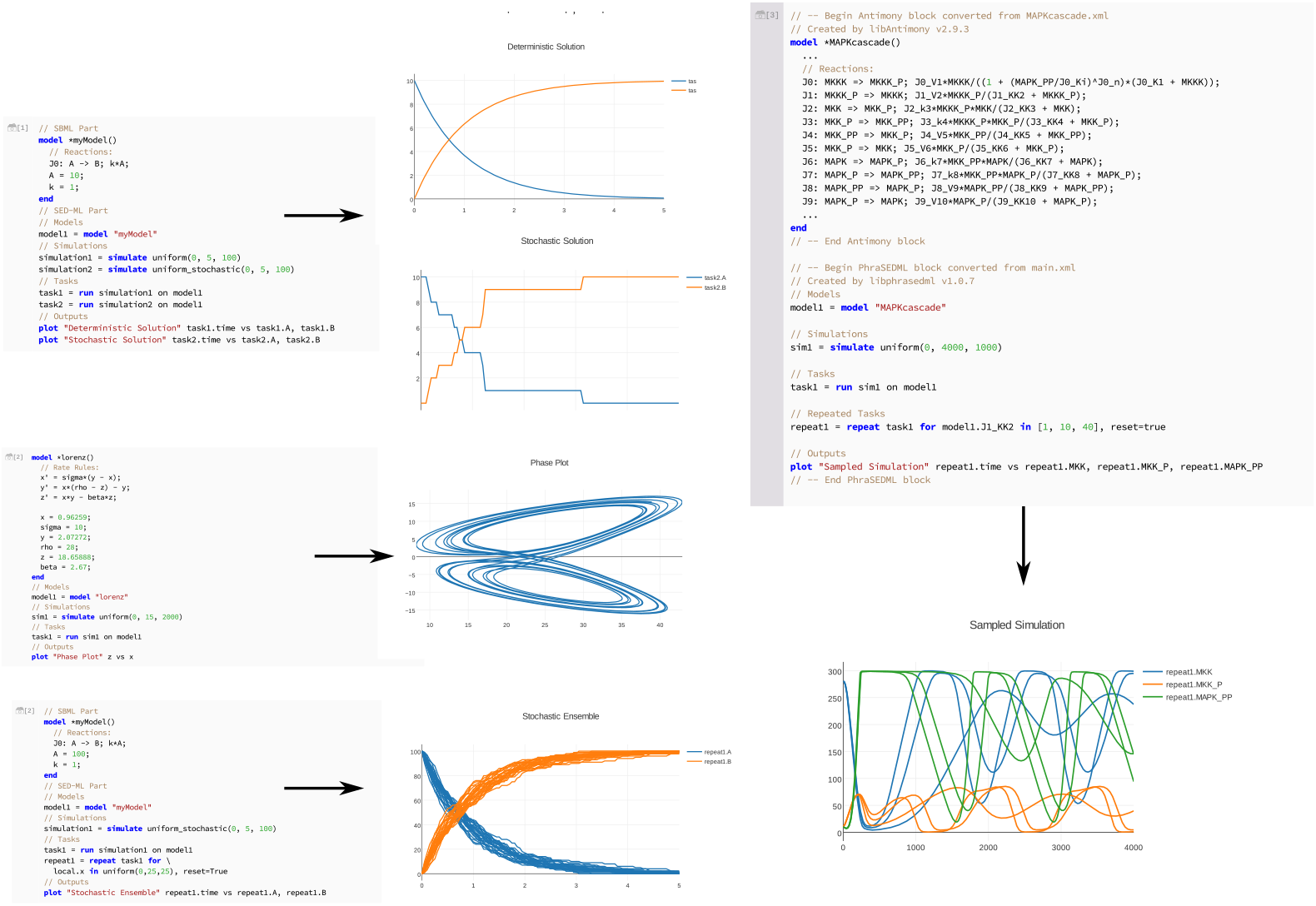
Examples of Tellurium’s inline OMEX format for specifying COMBINE archives. These examples start with very simple cases and build on these cases with progressively more advanced features. The first example contains a simple two–species model and simulated using a deterministic and stochastic solver. The second example shows a phase plot. The third example shows multiple stochastic traces. The fourth example shows a one–dimensional parameter scan. All of these examples are available via a Tellurium notebook, which can be accessed by clicking on “File” → “Open Example Notebook” → “COMBINE Archive Basics” from within the Tellurium notebook viewer.

**Fig S5.**
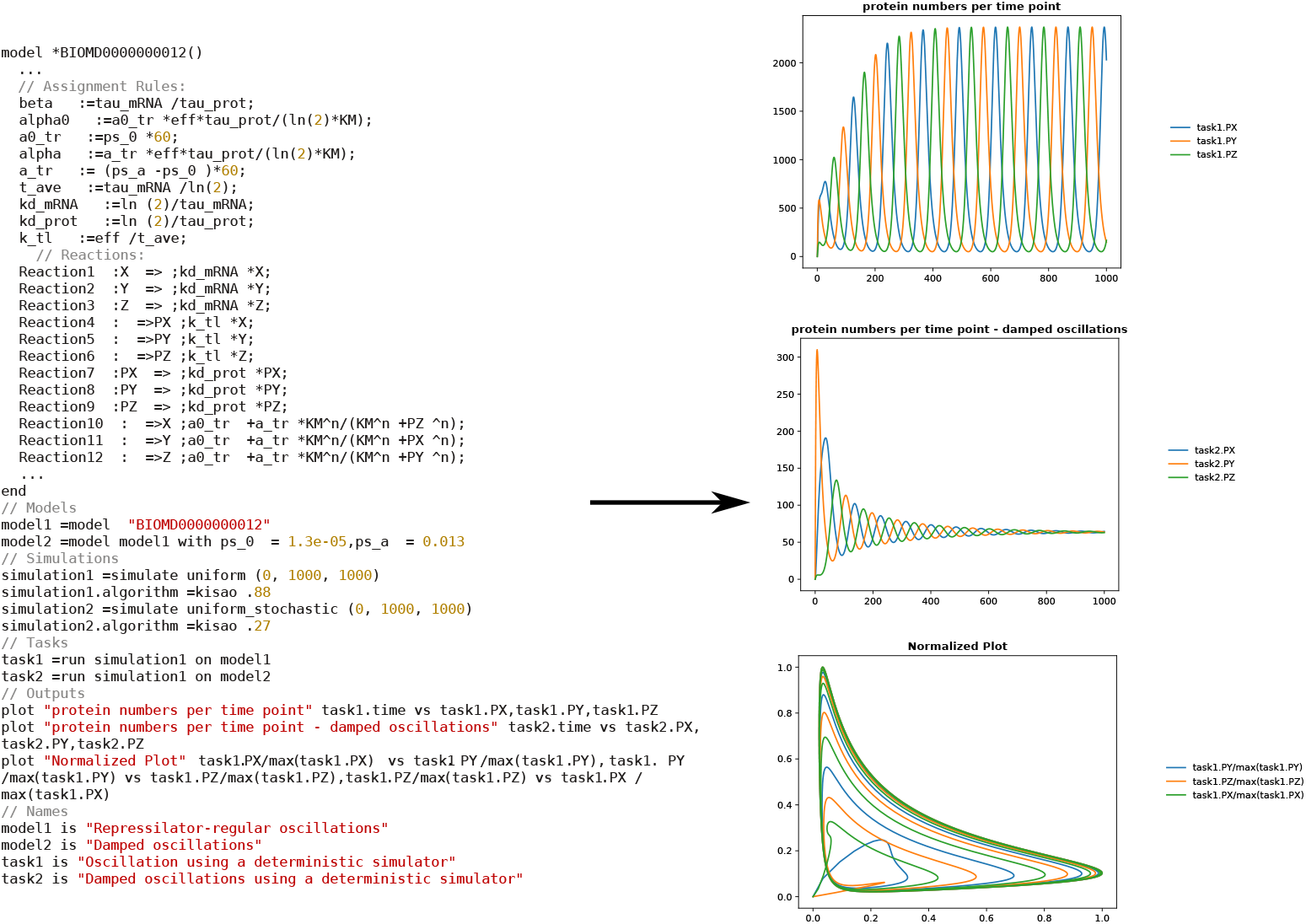
A normative example of a COMBINE archive introduced in the original paper describing the COMBINE archive format [21]. This example contains the repressilator model [79], a damped oscillation variant showing the modification of model parameters using SED–ML, and a phase plot of the undamped system.

**Table S1.** Combine Archive Test Cases.

|  |
| --- |
| **demos:** A set of models and SED-ML simulations of increasing complexity, in order to facilitate incremental support of the standard (4 archives, shown in Fig S4). |
| **real-models:** A selection of BioModels converted to COMBINE archives containing the associated SED-ML simulations (2 archives). |
| **published:** COMBINE archives published in the literature, including normative examples from the original paper introducing COMBINE archives [21] shown in Fig S5 (2 archives). |
| **swt:** Examples from the SED-ML Web Tools [22] demonstrating advanced features of SED-ML (5 archives). |
| **sbml-test-suite:** COMBINE archive encodings of the entire SBML test suite (1196 archives). |

We collected all COMBINE archives used during development of Tellurium in an online repository on GitHub [80]. These archives serve to test Tellurium’s standards compliance, but they may also allow other tool developers to better support COMBINE archives. We have therefore organized the test cases into different categories, from toy examples using progressively more advanced features of SED–ML, to BioModels, and finally advanced SED–ML usage. The test suite draws archives from a wide range of sources: publications [21,81], other tools (e.g. the SED–ML Web Tools [22]), the SBML Test Suite encoded as COMBINE archives, and archives developed by our group. The COMBINE test suite contains archives ranging from basic examples to advanced usage of the SED–ML standard. To verify exchangeability, we have manually tested importing these archives into our software and also into the SED–ML Web Tools. The SBML test cases were too numerous to test in this way, so a subset of archives were tested with the SED–ML Web Tools whereas the full set of archives was tested with Tellurium using a Tellurium notebook [82].

**Table S1.**
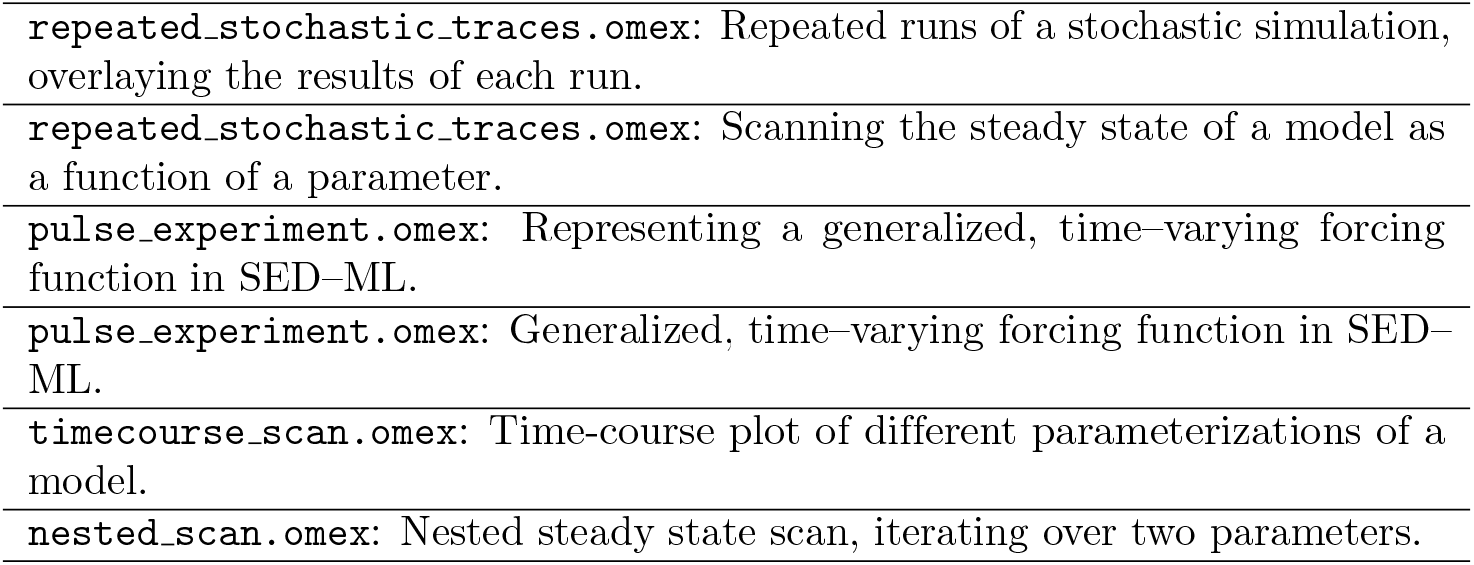
Advanced SED—ML Tests (Provided by SED—ML Web Tools [22]).

The SBML test suite was converted into COMBINE archives using the provided notebook https://github.com/0u812/tellurium-combine-archive-test-cases/blob/master/sbml-test-suite/convert-to-combine-arch.ipynb. These SBML test cases are automatically converted into COMBINE archives containing the expected results, which are then converted by Tellurium into inline OMEX and simulated. A “failed” test refers to a case where the numeric simulation results diverge from the expected values. An “unsupported” test refers to a test that uses features not available in our simulator (libroadrunner) or the inline OMEX strings.

**Table S3.**
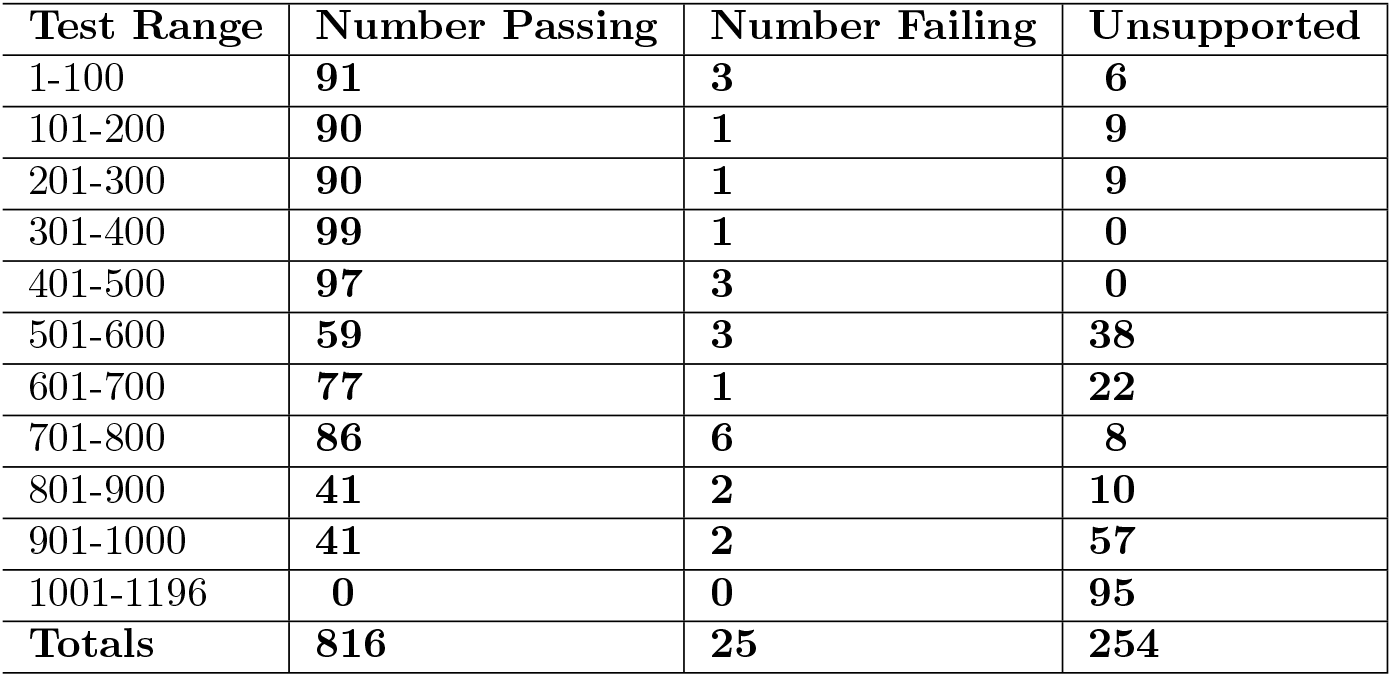
Tellurium SBML Test Suite Results.

The SBML test suite was converted into COMBINE archives using the provided notebook https://github.com/0u812/tellurium-combine-archive-test-cases/blob/master/sbml-test-suite/convert-to-combine-arch.ipynb. These SBML test cases are automatically converted into COMBINE archives containing the expected results, which are then converted by Tellurium into inline OMEX and simulated. A “failed” test refers to a case where the numeric simulation results diverge from the expected values. An “unsupported” test refers to a test that uses features not available in our simulator (libroadrunner) or the inline OMEX strings.

**Fig S6.**
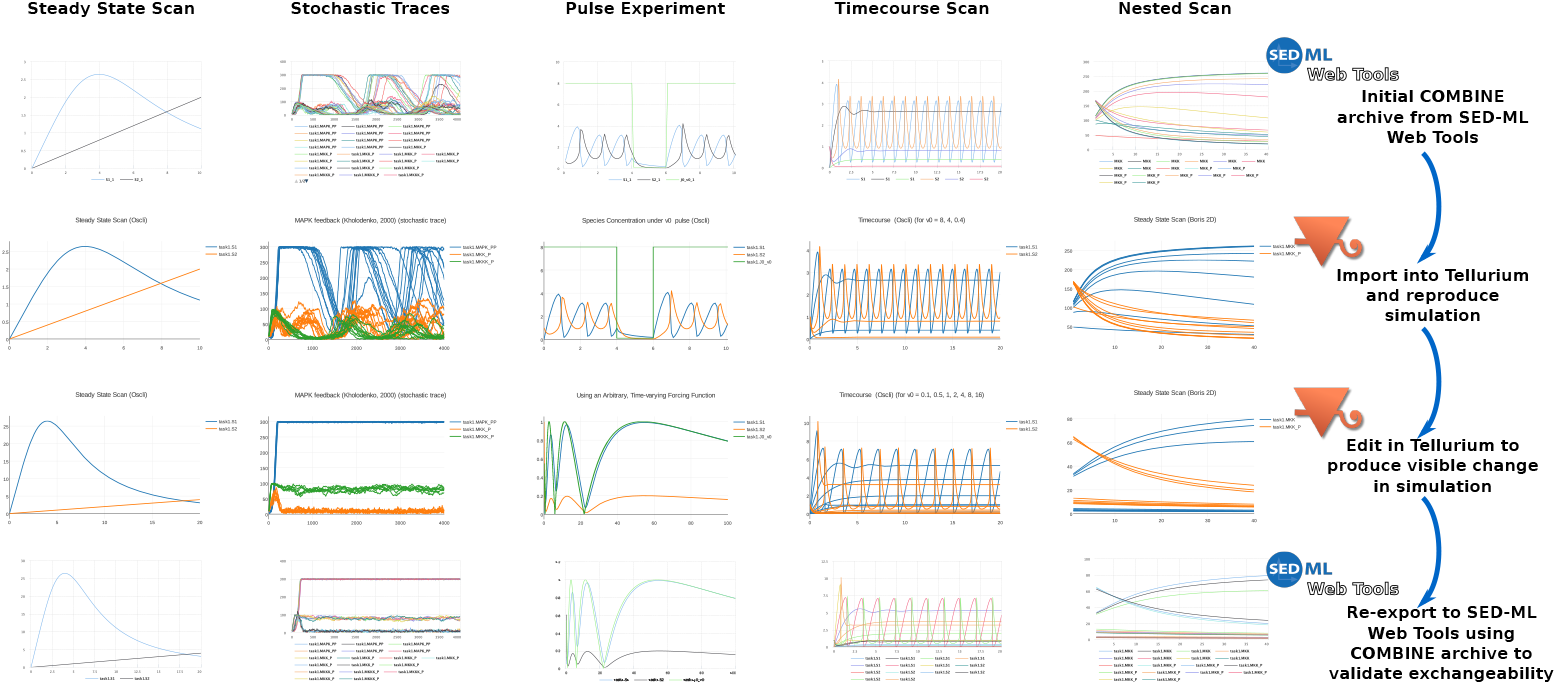
Round-tripping the SED–ML Web Tools examples [22]. In order to demonstrate broad support for standards, we conducted a series of tests utilizing advanced usage of SED–ML. The first row shows the original example rendered in the SED–ML Web Tools. The second row shows the same example imported into Tellurium. The third row shows the simulation after editing the model in Tellurium. Finally, the fourth row shows the result of re–exporting the example to the SED–ML Web Tools using a COMBINE archive.

## Footnotes

1 https://www.ebi.ac.uk/biomodels-main/BIOMD0000000144

## References

1 Karr JR, Sanghvi JC, Macklin DN, Gutschow MV, Jacobs JM, Bolival B, et al. A whole-cell computational model predicts phenotype from genotype. Cell. 2012;150(2):389–401.

2 Millard P, Smallbone K, Mendes P. Metabolic regulation is sufficient for global and robust coordination of glucose uptake, catabolism, energy production and growth in Escherichia coli. PLoS computational biology. 2017;13(2):e1005396.

3 Prinz F, Schlange T, Asadullah K. Believe it or not: how much can we rely on published data on potential drug targets? Nature reviews Drug discovery. 2011;10(9):712–712.

4 Mobley A, Linder SK, Braeuer R, Ellis LM, Zwelling L. A survey on data reproducibility in cancer research provides insights into our limited ability to translate findings from the laboratory to the clinic. PLoS One. 2013;8(5):e63221.

5 Peng GC. Moving Toward Model Reproducibility and Reusability. IEEE Transactions on Biomedical Engineering. 2016;63(10):1997–1998.

6 Medley JK, Goldberg AP, Karr JR. Guidelines for reproducibly building and simulating systems biology models. IEEE Transactions on Biomedical Engineering. 2016;63(10):2015–2020.

7 McDougal RA, Bulanova AS, Lytton WW. Reproducibility in computational neuroscience models and simulations. IEEE Transactions on Biomedical Engineering. 2016;63(10):2021–2035.

8 Waltemath D, Wolkenhauer O. How modeling standards, software, and initiatives support reproducibility in systems biology and systems medicine. IEEE Transactions on Biomedical Engineering. 2016;63(10):1999–2006.

9 Peng RD. Reproducible research in computational science. Science. 2011;334(6060):1226–1227.

10 Sandve GK, Nekrutenko A, Taylor J, Hovig E. Ten simple rules for reproducible computational research. PLoS computational biology. 2013;9(10):e1003285.

11 Hucka M, Finney A, Sauro HM, Bolouri H, Doyle JC, Kitano H, et al. The systems biology markup language (SBML): a medium for representation and exchange of biochemical network models. Bioinformatics. 2003;19(4):524–531.

12 Cuellar AA, Lloyd CM, Nielsen PF, Bullivant DP, Nickerson DP, Hunter PJ. An overview of CellML 1.1, a biological model description language. Simulation. 2003;79(12):740–747.

13 SBML Flux Balance Constraints.;. http://sbml.org/Documents/Specifications/SBML_Level_3/Packages/Flux_Balance_Constraints_(flux).

14 Waltemath D, Adams R, Bergmann FT, Hucka M, Kolpakov F, Miller AK, et al. Reproducible computational biology experiments with SED-ML-the simulation experiment description markup language. BMC systems biology. 2011;5(1):198.

15 Courtot M, Juty N, Knüpfer C, Waltemath D, Zhukova A, Dräger A, et al. Controlled vocabularies and semantics in systems biology. Molecular systems biology. 2011;7(1):543.

16 Petzold L. Automatic selection of methods for solving stiff and nonstiff systems of ordinary differential equations. SIAM journal on scientific and statistical computing. 1983;4(1):136–148.

17 Hindmarsh AC. ODEPACK, a systematized collection of ODE solvers;.

18 Cohen SD, Hindmarsh AC, Dubois PF, et al. CVODE, a stiff/nonstiff ODE solver in C. Computers in physics. 1996;10(2):138–143.

19 Gillespie DT. Exact stochastic simulation of coupled chemical reactions. The journal of physical chemistry. 1977;81(25):2340–2361.

20 Gibson MA, Bruck J. Efficient exact stochastic simulation of chemical systems with many species and many channels. The journal of physical chemistry A. 2000;104(9):1876–1889.

21 Bergmann FT, Adams R, Moodie S, Cooper J, Glont M, Golebiewski M, et al. COMBINE archive and OMEX format: one file to share all information to reproduce a modeling project. BMC bioinformatics. 2014;15(1):369.

22 Bergmann FT, Nickerson D, Waltemath D, Scharm M. SED-ML web tools: generate, modify and export standard-compliant simulation studies. Bioinformatics. 2017;33(8):1253–1254.

23 Scharm M, Wendland F, Peters M, Wolfien M, Theile T, Waltemath D. The CombineArchiveWeb application–A web based tool to handle files associated with modelling results. PeerJ PrePrints; 2014.

24 Ragan-Kelley M, Perez F, Granger B, Kluyver T, Ivanov P, Frederic J, et al. The Jupyter/IPython architecture: a unified view of computational research, from interactive exploration to communication and publication. In: AGU Fall Meeting Abstracts; 2014.

25 Wolfram S. Mathematica. Cambridge university press Cambridge; 1996.

26 Smith LP, Bergmann FT, Chandran D, Sauro HM. Antimony: a modular model definition language. Bioinformatics. 2009;25(18):2452–2454.

27 Choi K, Smith LP, Medley JK, Sauro HM. phraSED-ML: A paraphrased, human-readable adaptation of SED-ML. Journal of Bioinformatics and Computational Biology. 2016; p. 1650035.

28 Wolf J, Sohn HY, Heinrich R, Kuriyama H. Mathematical analysis of a mechanism for autonomous metabolic oscillations in continuous culture of Saccharomyces cerevisiae. FEBS letters. 2001;499(3):230–234.

29 Murray DB, Beckmann M, Kitano H. Regulation of yeast oscillatory dynamics. Proceedings of the National Academy of Sciences. 2007;104(7):2241–2246.

30 Winfree AT. Biological rhythms and the behavior of populations of coupled oscillators. Journal of theoretical biology. 1967;16(1):15–42.

31 Dodd AN, Salathia N, Hall A, Kévei E, Tóth R, Nagy F, et al. Plant circadian clocks increase photosynthesis, growth, survival, and competitive advantage. Science. 2005;309(5734):630–633.

32 BIOMD0000000090.;. https://www.ebi.ac.uk/biomodels-main/BIOMD0000000090.

33 Wolf 2001 Oxygen–only Plot.;. https://github.com/0u812/tellurium-combine-archive-test-cases/blob/master/biomodels/wolf2001-oxy-only.omex.

34 Petzold L, Hindmarsh A. Lsoda. Computing and Mathematics Research Division, I-316 Lawrence Livermore National Laboratory, Livermore, CA. 1997;94550.

35 Myers CJ, Barker N, Jones K, Kuwahara H, Madsen C, Nguyen NPD. iBioSim: a tool for the analysis and design of genetic circuits. Bioinformatics. 2009;25(21):2848–2849.

36 Calzone L, Thieffry D, Tyson JJ, Novak B. Dynamical modeling of syncytial mitotic cycles in Drosophila embryos. Molecular systems biology. 2007;3(1):131.

37 Gilbert S. Developmental Biology. 6th ed. Sinauer Associates; 2000. Available from: https://www.ncbi.nlm.nih.gov/books/NBK10081/.

38 Edgar BA, Sprenger F, Duronio RJ, Leopold P, O’Farrell PH. Distinct molecular mechanism regulate cell cycle timing at successive stages of Drosophila embryogenesis. Genes & development. 1994;8(4):440–452.

39 Huang Jy, Raff JW. The dynamic localisation of the Drosophila APC/C: evidence for the existence of multiple complexes that perform distinct functions and are differentially localised. Journal of cell science. 2002;115(14):2847–2856.

40 Raff JW, Jeffers K, Huang Jy. The roles of Fzy/Cdc20 and Fzr/Cdh1 in regulating the destruction of cyclin B in space and time. The Journal of cell biology. 2002;157(7):1139–1149.

41 martin scharm, Touré V. COMBINE Archive Show Case. 2016;doi:10.6084/m9.figshare.3427271.v1.

42 Calzone L, Fages F, Soliman S. BIOCHAM: an environment for modeling biological systems and formalizing experimental knowledge. Bioinformatics. 2006;22(14):1805–1807.

43 Reproduction of Calzone et al. Fig 1 and 3 as a COMBINE Archive.;. https://github.com/0u812/tellurium-combine-archive-test-cases/blob/master/demos/calzone-fig1-fig3.omex.

44 Testing Feedback Regulation in the Calzone Model.;. https://github.com/0u812/tellurium-combine-archive-test-cases/blob/master/demos/calzone-feedback-study.omex.

45 SBML Test Suite.;. http://sbml.org/Software/SBML_Test_Suite.

46 Briat C, Gupta A, Khammash M. Antithetic integral feedback ensures robust perfect adaptation in noisy biomolecular networks. Cell systems. 2016;2(1):15–26.

47 Le Novere N, Bornstein B, Broicher A, Courtot M, Donizelli M, Dharuri H, et al. BioModels Database: a free, centralized database of curated, published, quantitative kinetic models of biochemical and cellular systems. Nucleic acids research. 2006;34(suppl 1):D689–D691.

48 Li C, Donizelli M, Rodriguez N, Dharuri H, Endler L, Chelliah V, et al. BioModels Database: An enhanced, curated and annotated resource for published quantitative kinetic models. BMC systems biology. 2010;4(1):92.

49 Olivier BG, Snoep JL. Web-based kinetic modelling using JWS Online. Bioinformatics. 2004;20(13):2143–2144.

50 Lloyd CM, Lawson JR, Hunter PJ, Nielsen PF. The CellML model repository. Bioinformatics. 2008;24(18):2122–2123.

51 Hoops S, Sahle S, Gauges R, Lee C, Pahle J, Simus N, et al. COPASI—a complex pathway simulator. Bioinformatics. 2006;22(24):3067–3074.

52 Mendes P, Hoops S, Sahle S, Gauges R, Dada J, Kummer U. Computational modeling of biochemical networks using COPASI. Methods Mol Biol. 2009; p. 17–59. doi:10.1007/978-1-59745-525-1_2.

53 Bergmann FT, Sauro HM. SBW-a modular framework for systems biology. In: Simulation Conference, 2006. WSC 06. Proceedings of the Winter. IEEE; 2006. p. 1637–1645.

54 PathwayDesigner;. http://pathwaydesigner.org/.

55 Funahashi A, Matsuoka Y, Jouraku A, Morohashi M, Kikuchi N, Kitano H. CellDesigner 3.5: A Versatile Modeling Tool for Biochemical Networks. Proc IEEE. 2008;96(8):1254–1265. doi:10.1109/JPROC.2008.925458.

56 Funahashi A, Morohashi M, Kitano H, Tanimura N. CellDesigner: a process diagram editor for gene-regulatory and biochemical networks. BIOSILICO. 2003;1(5):159 – 162. doi:http://dx.doi.org/10.1016/S1478-5382(03)02370-9.

57 Moraru II, Schaff JC, Slepchenko BM, Blinov M, Morgan F, Lakshminarayana A, et al. Virtual Cell modelling and simulation software environment. IET Syst Biol. 2008;2(5):352–362.

58 Schaff JC, Vasilescu D, Moraru II, Loew LM, Blinov ML. Rule-based modeling with Virtual Cell. Bioinformatics. 2016;32(18):2880–2882.

59 Blinov ML, Schaff JC, Vasilescu D, Moraru II, Bloom JE, Loew LM. Compartmental and Spatial Rule-Based Modeling with Virtual Cell. Biophysical Journal. 2017;113(7):1365–1372. doi:10.1016/j.bpj.2017.08.022.

60 Swat MH, Thomas GL, Belmonte JM, Shirinifard A, Hmeljak D, Glazier JA. Multi-scale modeling of tissues using CompuCell3D. Methods in cell biology. 2012;110:325.

61 Olivier BG, Rohwer JM, Hofmeyr JHS. Modelling cellular systems with PySCeS. Bioinformatics. 2005;21(4):560–561.

62 Blinov ML, Faeder JR, Goldstein B, Hlavacek WS. BioNetGen: software for rule-based modeling of signal transduction based on the interactions of molecular domains. Bioinformatics. 2004;20(17):3289–3291.

63 Lopez CF, Muhlich JL, Bachman JA, Sorger PK. Programming biological models in Python using PySB. Molecular systems biology. 2013;9(1):646.

64 Ebrahim A, Lerman JA, Palsson BO, Hyduke DR. COBRApy: constraints-based reconstruction and analysis for python. BMC systems biology. 2013;7(1):74.

65 Fan S, Geissmann Q, Lakatos E, Lukauskas S, Ale A, Babtie AC, et al. MEANS: python package for Moment Expansion Approximation, iNference and Simulation. Bioinformatics. 2016;32(18):2863–2865.

66 Swaminathan A, Hsiao V, Murray RM. Quantitative Modeling of Integrase Dynamics Using a Novel Python Toolbox for Parameter Inference in Synthetic Biology. bioRxiv. 2017;doi:10.1101/121152.

67 Liepe J, Barnes C, Cule E, Erguler K, Kirk P, Toni T, et al. ABC-SysBio—approximate Bayesian computation in Python with GPU support. Bioinformatics. 2010;26(14):1797–1799.

68 Theisen M. Introducing py_emra: the Python module for Ensemble Modeling Robustness Analysis. bioRxiv. 2016;doi:10.1101/065177.

69 Goecks J, Nekrutenko A, Taylor J. Galaxy: a comprehensive approach for supporting accessible, reproducible, and transparent computational research in the life sciences. Genome biology. 2010;11(8):R86.

70 Pond SK, Wadhawan S, Chiaromonte F, Ananda G, Chung WY, Taylor J, et al. Windshield splatter analysis with the Galaxy metagenomic pipeline. Genome research. 2009;19(11):2144–2153.

71 Oinn T, Addis M, Ferris J, Marvin D, Senger M, Greenwood M, et al. Taverna: a tool for the composition and enactment of bioinformatics workflows. Bioinformatics. 2004;20(17):3045–3054.

72 Callahan SP, Freire J, Santos E, Scheidegger CE, Silva CT, Vo HT. VisTrails: visualization meets data management. In: Proceedings of the 2006 ACM SIGMOD international conference on Management of data. ACM; 2006. p. 745–747.

73 Piccolo SR, Frampton MB. Tools and techniques for computational reproducibility. GigaScience. 2016;5(1):30.

74 Drawert B, Hellander A, Bales B, Banerjee D, Bellesia G, Daigle Jr BJ, et al. Stochastic Simulation Service: Bridging the Gap between the Computational Expert and the Biologist. PLoS computational biology. 2016;12(12):e1005220.

75 Reich M, Tabor T, Liefeld T, Thorvaldsdóttir H, Hill B, Tamayo P, et al. The GenePattern Notebook Environment. Cell Systems. 2017;5(2):149–151.e1. doi:10.1016/j.cels.2017.07.003.

76 Eröcal B, Stein W. The Sage Project: Unifying Free Mathematical Software to Create. In: Mathematical Software-ICMS 2010: Third International Congress on Mathematical Software, Kobe, Japan, September 13–17, 2010, Proceedings. vol. 6327. Springer; 2010. p. 12.

77 Ewald R, Uhrmacher AM. SESSL: A domain-specific language for simulation experiments. ACM Transactions on Modeling and Computer Simulation (TOMACS). 2014;24(2):11.

78 Hill Coefficient Study Based On Wolf 2001.;. https://github.com/0u812/tellurium-combine-archive-test-cases/blob/master/demos/wolf-hill-study.omex.

79 Elowitz MB, Leibler S. A synthetic oscillatory network of transcriptional regulators. Nature. 2000;403(6767):335–338.

80 Tellurium COMBINE Archive Tests.;. https://github.com/0u812/tellurium-combine-archive-test-cases.

81 Scharm M, Waltemath D. A fully featured COMBINE archive of a simulation study on syncytial mitotic cycles in Drosophila embryos. F1000Research. 2016;.

82 Tellurium SBML Test Suite Notebook.;. https://github.com/0u812/tellurium-combine-archive-test-cases/blob/master/sbml-test-suite/convert-to-combine-arch.ipynb.

